# Zn^2+^ potentiation of OTOP proton channels identifies structural elements of the gating apparatus

**DOI:** 10.1101/2022.12.15.520539

**Authors:** Bochuan Teng, Joshua P. Kaplan, Ziyu Liang, Marcel Goldschen-Ohm, Emily R. Liman

## Abstract

Otopetrin proteins (OTOPs) form proton-selective ion channels that are expressed in diverse cell types where they may mediate detection of acids or regulation of pH. In vertebrates there are three family members: OTOP1 is required for formation of otoconia in the vesibular system and it forms the receptor for sour taste, while the functions of OTOP2, and OTOP3 are not yet known. Importantly, the gating mechanisms of any of the OTOP channels are not well-understood, and until recently, it was not even known if the channels were gated. Here we show that Zn^2+^, as well as other transition metals including copper (Cu^2+^), potently activate murine OTOP3. Zn^2+^ pre-exposure increases the magnitude of OTOP3 currents to a subsequent acid stimulus by as much as 10-fold. In contrast, OTOP2 currents are insensitive to potentiation by Zn^2+^. Swapping the extracellular tm 11-12 linker between OTOP3 and OTOP2 was sufficient to eliminate Zn^2+^ potentiation of OTOP3 and confer Zn^2+^ potentiation on OTOP2. We also show that H531 within the tm 11-12 linker is essential for potentiation of OTOP3 by Zn^2+^, likely by forming part of its binding site. Kinetic modeling of the data is consistent with Zn^2+^ stabilizing the open state of the channel, possibly competing with H^+^ for activation of the channels. These results establish the tm 11-12 linker as part of the gating apparatus of OTOP channels and a target for drug discovery. Zinc is an essential micronutrient and its regulation of OTOP channels will undoubtedly have important physiological sequelae.

**Significance Statement:** A family of proton-activated H^+^ ion channels was recently identified that includes the sour receptor OTOP1. Here we show that members of the OTOP channel family are differentially sensitive to Zn^2+^, which strongly activates OTOP3 but not OTOP2. By studying chimeric channels, we identify structural elements in the extracellular linker between transmembrane domains 11-12 that are necessary and sufficient for Zn^2+^ activation of OTOP channels. In addition to the taste system, OTOP channels play important roles in biomineralization in both vertebrate and invertebrates and are expressed in the digestive tract where their expression is a predictor of cancer prognosis. Our identification of the tm 11-12 linker as part of the gating apparatus makes it a promising target for pharmaceutical discovery.

## Introduction

Pharmacological agents that can activate or inhibit ion channels have long been used as probes to describe the fundamental processes of channel gating and ion permeation (Hille, 2001). For example, the discovery of the charged molecule TEA and the scorpion toxin charybdotoxin as a specific blocker of K^+^ channels allowed for the early identification of residues lining the channel pore well before the channel structures were determined (MacKinnon et al., 1990; Yellen et al., 1991; Banerjee et al., 2013). Similarly, gating modifiers have been used to probe structural rearrangements that accompany the opening of voltage-gated ion channels (Swartz and MacKinnon, 1997; Sack and Aldrich, 2006; Catterall et al., 2007; Goldschen-Ohm and Chanda, 2014). More recently, toxins that target pain-sensing ASIC and TRPV1 channels have been used to probe the conformational states of these channels (Bohlen et al., 2010; Baconguis et al., 2014). One of the most common modulators of channel activity is the trace metal zinc (Zn^2+^), which can affect gating, permeation, or both (Gilly and Armstrong, 1982; Chu et al., 2004; Noh et al., 2015; Peralta and Huidobro-Toro, 2016). Zn^2+^ binds to proteins with high affinity and specificity and regulates a wide range of cellular processes, including metabolism and gene expression (Vallee and Falchuk, 1993). Zn^2+^ is a potent inhibitor of proton-transport molecules including the voltage-gated proton channel Hv1 and the proton-selective ion channel OTOP1 (Decoursey, 2003; Ramsey et al., 2006; Tu et al., 2018).

OTOP1 is a member of a family of proteins (Hughes et al., 2008; Hurle et al., 2011), which includes, within vertebrates, two other members, OTOP2 and OTOP3, that also function as proton channels (Tu et al., 2018). OTOP proton channels are expressed throughout the body, where they play diverse and still poorly understood roles in pH sensing and homeostasis. In vertebrates and invertebrates, OTOP channels expressed in the gustatory system sense acids and function as sour taste receptors (OTOP1 for vertebrates; OTOPL1 for Drosophila) (Teng et al., 2019; Zhang et al., 2019; Ganguly et al., 2021; Mi et al., 2021). In mice and zebrafish, OTOP1 plays an essential role in the formation of force-sensing calcium carbonate-based otoconia in the ear (Hurle et al., 2003; Hughes et al., 2004), likely by regulating pH in the endolymph. OTOP2 and OTOP3 are both found throughout the digestive system, and their expression has been shown to correlate with disease progression in some forms of colon cancer (Tu et al., 2018; Parikh et al., 2019; Yang et al., 2019). Most recently, an OTOP channel was shown to be critically involved in calcification and the formation of a skeleton in sea urchin embryos (Chang et al., 2021).

Given the recent discovery of OTOP proteins as forming ion channels (Tu et al., 2018), much remains to be discovered about how they function. For example, it was not known if the channels occupy open and closed states or if those terms even apply to these proteins, which bear no structural similarity to other ion channels, and that could conduct protons through a non-aqueous pathway (Decoursey, 2003). Recently, we showed that OTOP channels are gated by extracellular protons, acting mostly likely on multiple titratable residues on the extracellular domain of the protein (Teng et al., 2022). Here we report the first evidence that OTOP channels can be potentiated by Zn^2+^ and Cu^2+^, with current magnitudes augmented by up to 10-fold. This is direct evidence that the OTOP channels are gated, like nearly all other ion channels. Using a chimeric channel approach, we identify key determinants for Zn^2+^ potentiation that likely represent elements of the gating apparatus of the channels and potential targets for pharmacological manipulation.

## Results

### Zn^2+^ both blocks and potentiates OTOP3 currents

While measuring the sensitivity of the three murine OTOP channels to inhibition by Zn^2+^, we noticed that mOTOP3 currents were larger following removal of Zn^2+^ than before its introduction. We refer to this effect as potentiation. For these and other experiments, OTOP channels were expressed in HEK-293 cells and studied by patch clamp recording. As shown in Figure 1, all three OTOP channels carried inward currents in response to a pH 5.5. stimulus and were subsequently inhibited by 1 mM Zn^2+^ at pH 5.5. However, only mOTOP3 currents showed recovery during the Zn^2+^ exposure and a large rebound, of nearly 3-fold, following its removal (Figure 1 A, B). The potentiating effect of Zn^2+^ was dose-dependent over a concentration range of 0.1 mM – 10 mM and did not show evidence of saturation (Figure 1C, D). This is in contrast to the inhibitory effect of Zn^2+^ on mOTOP3 currents, which for a stimulus of pH 5.5 could be fit with an IC50 = 0.31 mM (Figure 1D). The difference in the dose-dependence of inhibition and potentiation suggests that Zn^2+^ interacts with distinct binding sites on the channels to produce the two effects (see below).

**Figure 1.**
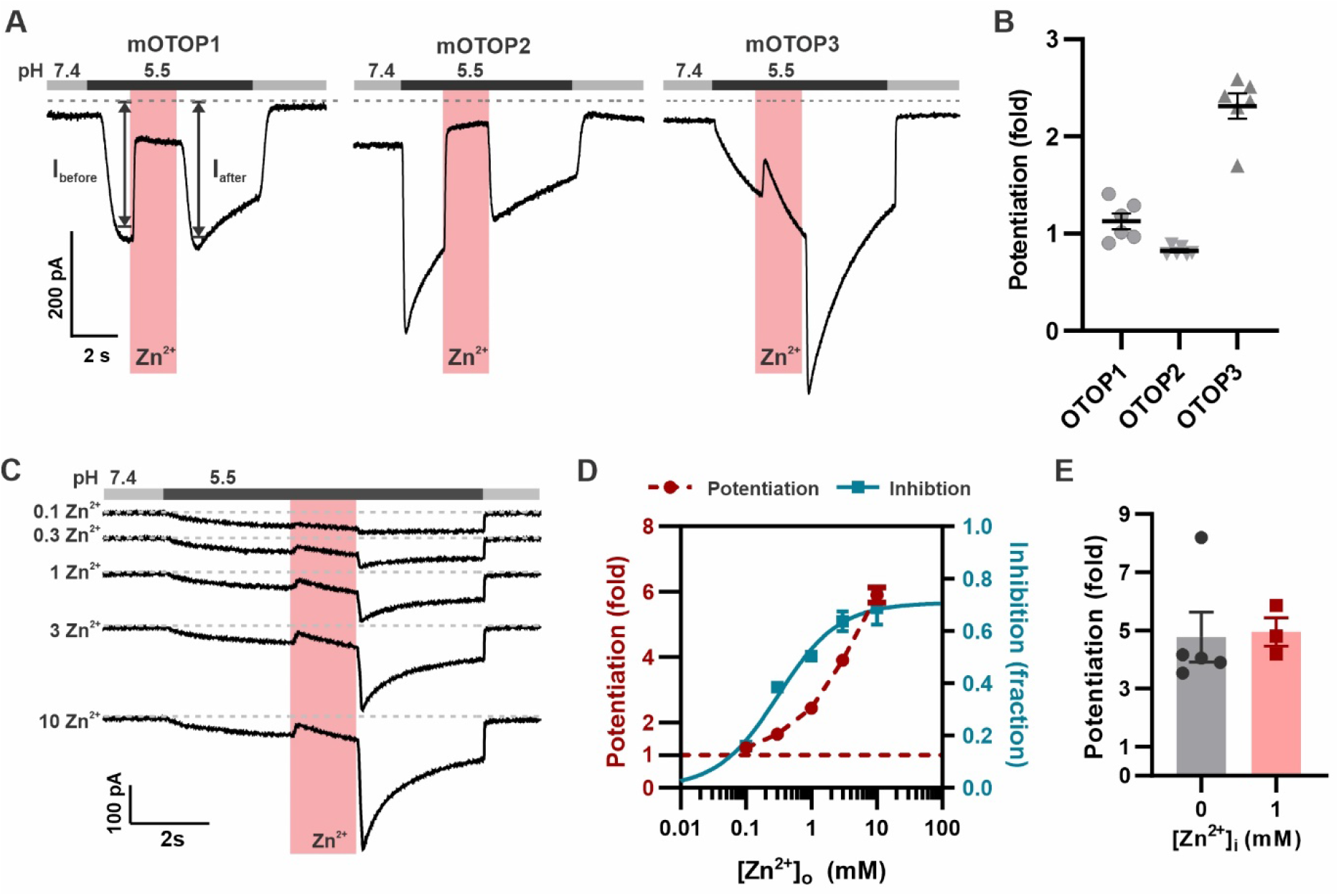
Zn^2+^ blocks and potentiates OTOP3 currents. **A.** Representative traces show Zn^2+^ both inhibits and potentiates mOTOP3 currents. Proton currents were elicited in HEK293 cells expressing each of the three OTOP channels in response to lowering the extracellular pH to 5.5 in the absence of extracellular Na^+^ as indicated. Zn^2+^ (pink bar, 1 mM) inhibits currents through all three channels, but only OTOP3 currents are potentiated following Zn^2+^ removal. Vm was held at −80 mV. **B.** Average and all points data for experiments as in **A** showing fold-potentiation as measured by comparing the current magnitude before and after Zn^2+^ application (arrows shown in **A**). **C.** Zn^2+^ applied at varying concentrations (pink bar, concentration indicated in mM) produces a dose-dependent inhibition and potentiation of OTOP3 currents. **D.** Average data from experiments in C show the dose-dependence of potentiation and inhibition (n=4 for Zn^2+^ potentiation, n=6 for Zn^2+^ inhibition). The dose-dependence of Zn^2+^ inhibition was fit with a Hill equation, with an IC_50_ = 0.31 mM, and Hill coefficient = 0.94. **E.** Average potentiation of OTOP3 currents in response to 1mM extracellular Zn^2+^ with (grey) or without (pink) 1mM Zn^2+^ loaded in the pipette. There was no difference between the two conditions (Student’s t-test, P>0.05).

We next asked whether the effect of Zn^2+^ to potentiate the mOTOP3 currents was through actions on the extracellular or the intracellular side of the channel, possibly following entry into the cytosol as is the case for TRPA1 (Hu et al., 2009). Introducing 1mM Zn^2+^ into the patch-pipette did not change the degree of potentiation in response to 1 mM extracellular Zn^2+^, indicating that Zn^2+^likely acts on extracellular domains of the channel (Figure 1E).

### Pre-exposure to Zn^2+^ potentiates OTOP3 currents

To study the activating effects of Zn^2+^ on mOTOP3 and avoid confounds due to its inhibitory effect, we devised a recording protocol in which the cells were pre-exposed to Zn^2+^ at pH 7.4, prior to evoking currents with an acidic stimulus. As shown in Figure 2A, this produced a robust potentiation of mOTOP3 currents evoked in response to the pH 5.5 stimulus. Notably, the currents were both faster and larger after exposure to Zn^2+^.

**Figure 2.**
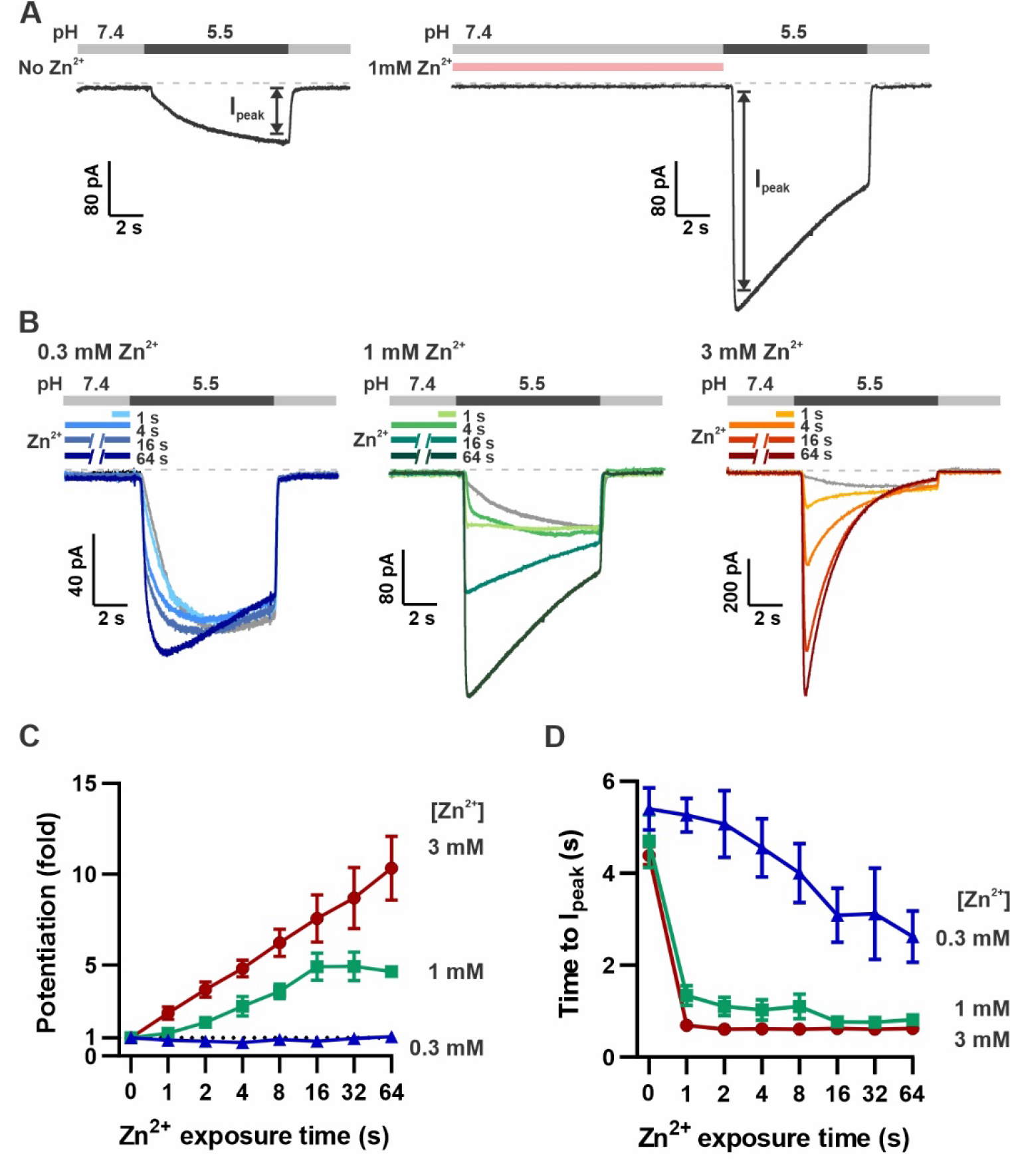
Pre-exposure to Zn^2+^ potentiates OTOP3 currents in a dose- and time-dependent manner. **A.** Solution exchange protocol designed to measure effects of Zn^2+^ on gating of OTOP currents without confounds due to its inhibitory effects. V_m_ was held at −80 mV. In this example, currents were elicited to a pH 5.5 stimulus without pre-exposure to Zn^2+^ and then following a 16 s exposure to 1 mM Zn^2+^. **B.** Representative traces show OTOP3 currents elicited in response to pH 5.5 stimulus with pre-exposure to 0.3 mM (blue), 1 mM (green) and 3mM (orange/red) Zn^2+^ for durations from 1-64s as indicated. **C, D**. The fold potentiation (C) and time to I_peak_ (D) as a function of Zn^2+^ pre-exposure time from experiments as in B (n=5–7). Fold potentiation was measured as the ratio of the current evoked to the pH 5.5. stimulus after Zn^2+^ to the control response in the absence of Zn^2+^. Data are plotted as mean +/− s.e.m.

We measured the dose and time-dependence of potentiation by Zn^2+^, using three concentrations of Zn^2+^: 0.3 mM, 1 mM, and 3 mM and by varying the duration of the Zn^2+^ pre-exposure from 1s – 64s. The response to Zn^2+^ was compared to the response in the absence of Zn^2+^ from the same cell. As shown in Figure 2 B, C., 3 mM Zn^2+^ caused a more than 10-fold increase in the peak current, the lowest concentration of Zn^2+^, 0.3 mM, applied for up to 64 seconds produced only a negligible increase in the peak current and 1 mM Zn^2+^ had an intermediate effect. Thus, the potentiating effect of Zn^2+^, as measured by the peak current, was dose-dependent, with an apparent threshold of > 0.3 mM Zn^2+^. Examination of the time-dependence of the response showed increasing potentiation with exposure times up to ~16 seconds for concentrations of 1 mM and 3 mM Zn^2+^, at which point the effect tended to saturate, although there was some variability from cell to cell (Figure 2C).

In addition to increasing the peak current, Zn^2+^ pre-exposure also increased the apparent rate of activation to a pH 5.5 stimulus that otherwise slowly activates mOTOP3 currents (Teng et al., 2022). This change in apparent activation kinetics showed a dose- and time-dependence (Figure 2D). At the higher concentrations (1 and 3 mM), an exposure of 1 second was sufficient to observe a maximal decrease in the time to peak current from ~ 4 seconds to < 1 second. At 0.3 mM Zn^2+^, this effect required longer exposure times, and activation rates never reached the speed obtained with 1-second exposure to 1 mM Zn^2+^.

To assess the stability of the Zn^2+^ bound conformation, we measured the rate of recovery from potentiation by Zn^2+^ (Figure 3). For these experiments, the cells were exposed to 1 mM Zn^2+^ (pH 7.4) for 16 s. This was followed by a “wash-off” period of 0-64 seconds in a Zn^2+^-free solution (pH 7.4) before currents were activated with a pH 5.5 solution (Figure 3A). A wash-off period of one second was sufficient to reduced potentiation by 50% while a period of >16s allowed for a complete recovery of currents to baseline.

**Figure 3.**
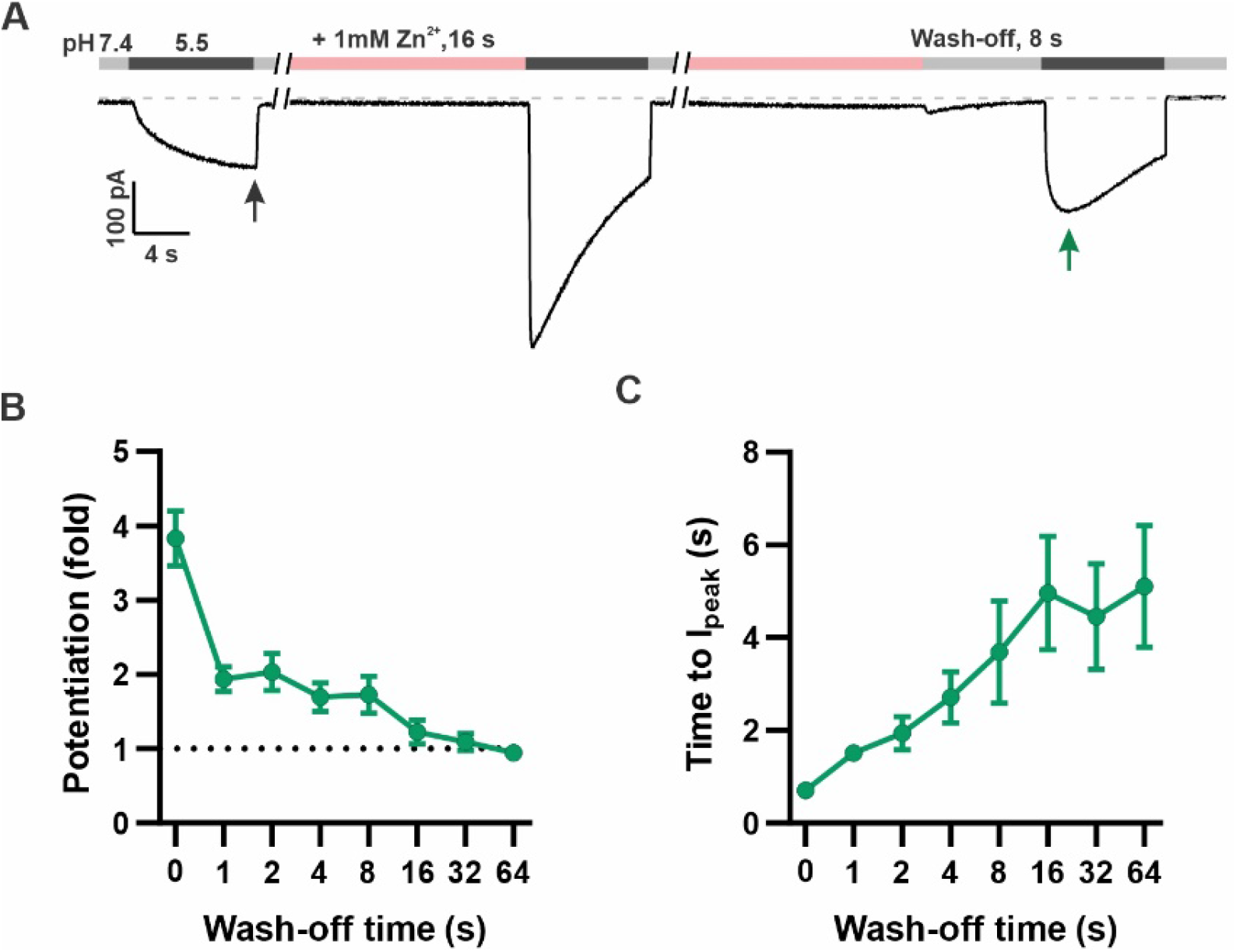
Time dependence of the recovery of OTOP3 currents from Zn^2+^ pre-potentiation. **A.** Solution exchange protocol designed to measure the recovery of OTOP3 currents following exposure to Zn^2+^. In this example, the cell expressing OTOP3 was first exposed to 1mM Zn^2+^ for 16s which was followed by an 8 s wash-off phase in pH 7.4 solution before currents were elicited in response to the pH 5.5 solution. **B, C**. The fold potentiation (B) and time to Ipeak (C) as a function of Zn^2+^ wash-off time from experiments as in A (n=4-6). Data are plotted as mean +/− s.e.m.

Thus, by applying Zn^2+^ at pH 7.4 before activation with a Zn^2+^-free solution at pH 5.5, we could elicit robust potentiation of mOTOP3 currents from which we could characterize the Zn^2+^ and time dependence of this effect, and the time-dependence of its recovery.

### OTOP1 is potentiated by Zn^2+^

Using this new protocol, we went back and assessed the effect of Zn^2+^ on mOTOP1 and mOTOP2. mOTOP1 and mOTOP2 currents evoked in response to a solution at pH 5.5 showed no evidence of potentiation by exposure to 1 mM Zn^2+^ applied for 16 seconds at pH 7.4 (Figure 4A, B). As mOTOP1 and mOTOP2 differ from mOTOP3 channels in that they display a greater degree of activation at pH 5.5 (Teng et al., 2022), we considered whether this might preclude further potentiation by Zn^2+^ (see Figure 8). Indeed, we found that mOTOP1 currents could be potentiated when activated by a milder stimulus, pH 6.0 by as much as 3-fold after a 64-second exposure to 1 mM Zn^2+^ (Figure 4C), with a time dependence similar to what we observed for mOTOP3 (Figure 4D). Activation rates of the more rapidly activating OTOP1 channels were not measurably enhanced by pre-exposure to Zn^2+^ exposure (Figure 4E). Thus, potentiation by Zn^2+^ is a feature shared by mOTOP1 and mOTOP3.

**Figure 4.**
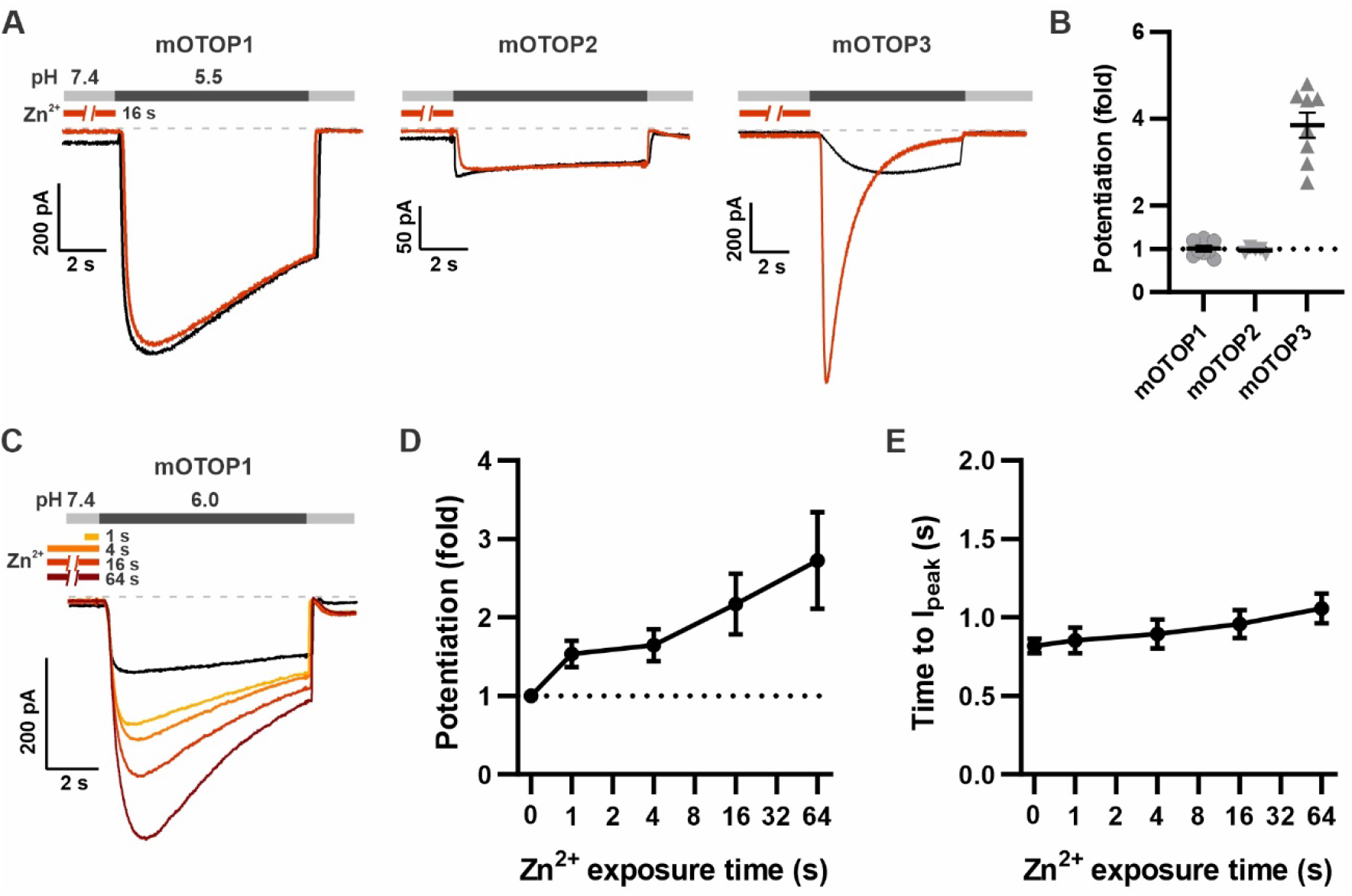
mOTOP1 is potentiated by Zn^2+^ when activated by a mild acid stimulus. **A.** Proton currents recorded from HEK293 cells expressing each of the three OTOP channels as indicated, in response to pH 5.5 with (red) or without (black) Zn^2+^ pre-exposure. V_m_ was held at −80 mV. The cells were exposed to 1mM Zn^2+^ for 16 s prior to pH 5.5 solutions. **B**. Average and all points data from experiments as in A showing the fold-potentiation in response to 1 mM Zn^2+^ for 16 s. **C.** Representative traces showing OTOP1 currents evoked in response to a pH 6.0 stimulus before and after exposure to Zn^2+^ for varying times as indicated. **D, E**. The fold potentiation (C) and time to I_peak_ (D) as a function of Zn^2+^ pre-exposure time from experiments as in C (n= 5). Data are plotted as mean +/− s.e.m.

### Divalent transition metal ions potentiate OTOP3

A wide range of ion channels and neurotransmitter receptors have been shown to be modulated by Zn^2+^, some of which are modulated by other divalent transition metals interacting with the same residues that bind Zn^2+^ (Mathie et al., 2006; Shcheglovitov et al., 2012). For example, the zinc-activated ion channel, a member of the family of Cys-loop receptors, is activated by copper and protons, as well as by zinc (Trattnig et al., 2016) while the cyclic-nucleotide gated ion channel from rods is potentiated by Ni^2+^, as well as by Zn^2+^, Cd^2+^ and Co^2+^, acting through a histidine residue in the mouth of the channel (Karpen et al., 1993; Gordon and Zagotta, 1995a). Divalent transition metals such as Co^2+^, Ni^2+^, Cu^2+^, and Cd^2+^ are predicted to have distinct preferred coordination geometries in metalloproteins and other proteins to which they bind but are often coordinated by the same acidic and/or polar residues including histidine, glutamic acid, aspartic acid, and cysteine (14). To gain insights into the nature of the Zn^2+^ binding site, we tested whether other transition metals could potentiate mOTOP3 currents. Each metal ion was presented at a concentration of 1 mM for 16 seconds prior to activation of currents with a pH 5.5 stimulus. Pre-exposure to Cu^2+^ caused a dramatic increase in the magnitudes of the currents, potentiating them by 11.6 +/− 0.1-fold (n=10). Pre-exposure to Cd^2+^ and Ni^2+^ had more modest effects, potentiating mOTOP3 currents by 1.7 +/− 0.1 (n=10) and 1.6-fold +/− 0.0 (n=8). In contrast, Co^2+^ and Fe^2+^ had little to no effect on the magnitude or kinetics of the currents (Figure 5). These results suggest that the binding site occupied by Zn^2+^ may also be shared by other d-block transition metals.

**Figure 5.**
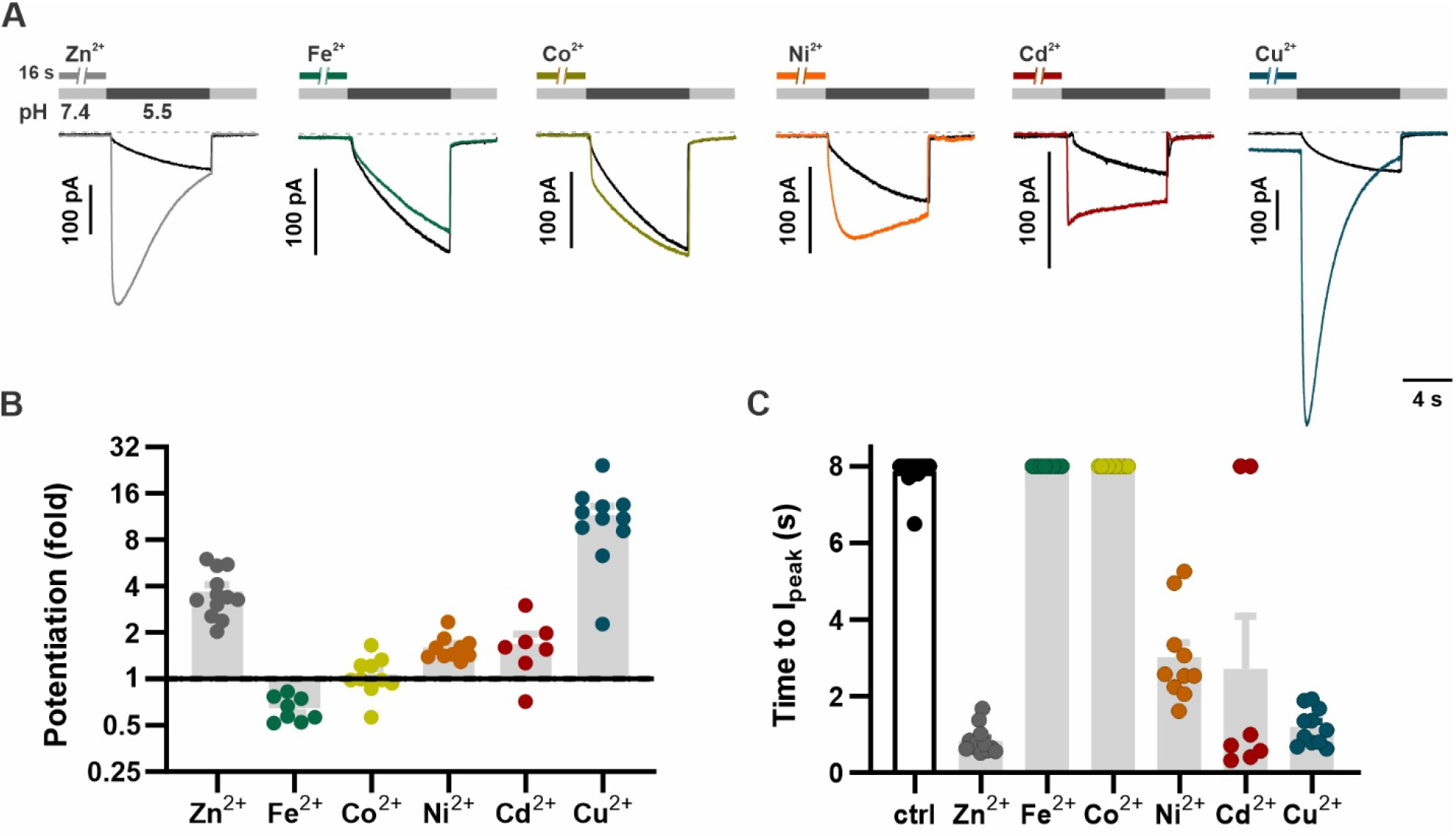
Divalent transition metal ions also potentiate OTOP3. **(A)** Proton currents in response to a pH 5.5 stimulus following exposure (1 mM, 16 s) to various d-block transition metals recorded from HEK293 cells expressing wildtype mOTOP3. V_m_ was held at −80 mV. Black trace is the control from the same experiment (cell). **(B)** Average (mean +/− s.e.m.) and all points data showing the fold-potentiation measured from experiments as in A. **(C)** Average (mean +/− s.e.m.) and all points data for latency to I_peak_, measured from experiments as in A. The latency to peak in C was scored as 8s when peak magnitudes were not reached before 8s.

### The tm 11-12 linker is necessary and sufficient to confer sensitivity to Zn^2+^ potentiation

Given the striking difference between mOTOP3 and mOTOP2 in potentiating effects of Zn^2+^, we reasoned that chimeras between the two channels might allow us to identify its structural basis. As Zn^2+^ is likely to bind to an extracellular domain, we tested chimeras in which each of the six external linkers between transmembrane domains were exchanged (Teng et al., 2022). A total of twelve chimeras were tested, six in which the backbone was the mOTOP2 channel and six in which the backbone was the mOTOP3 channels. Each chimera was tested for potentiation following pre-exposure to 1mM Zn^2+^ for 16s (Figure 6).

**Figure 6.**
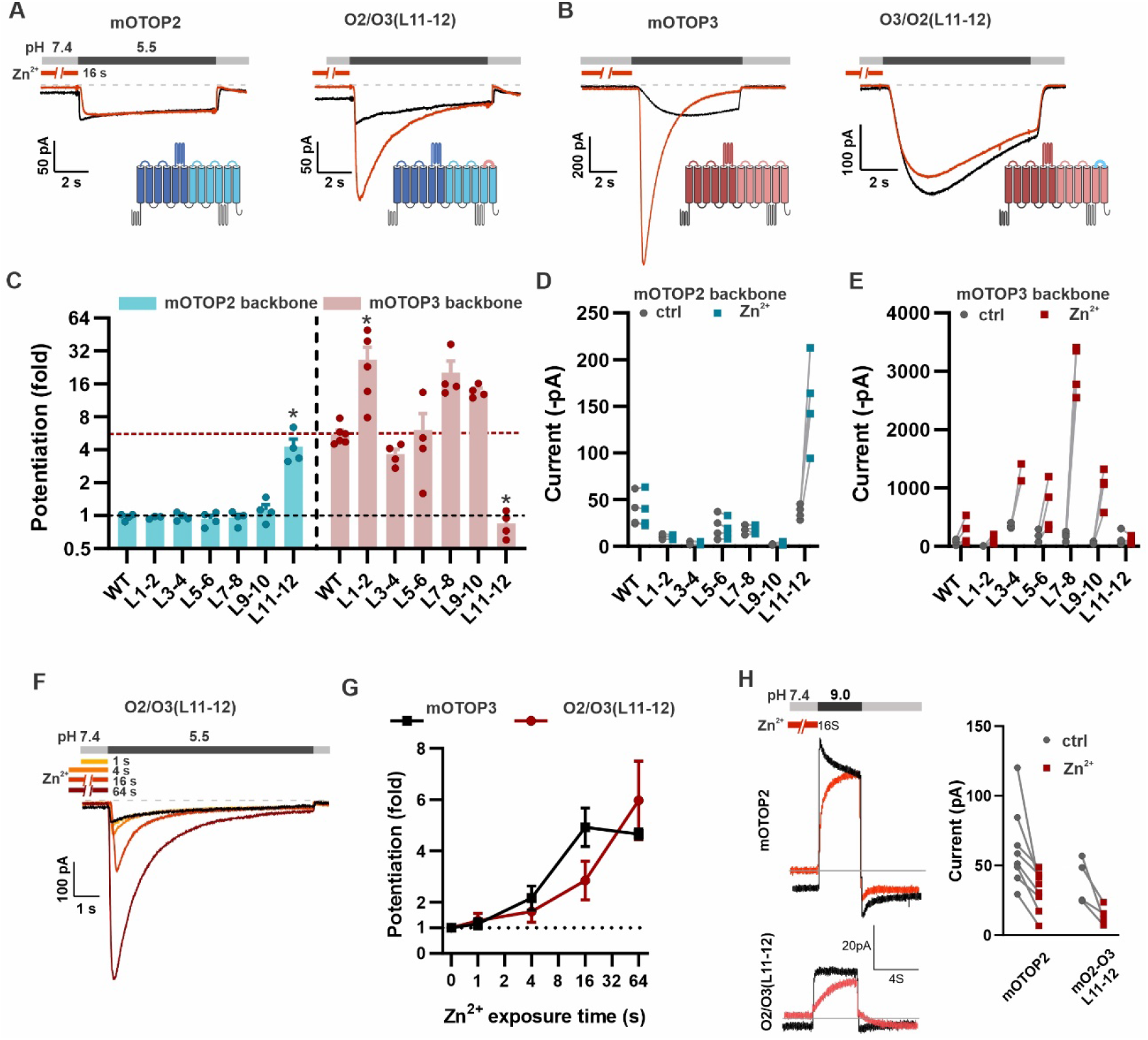
The tm 11-12 linker is both necessary and sufficient for Zn^2+^ potentiation. **A, B.** Proton currents in response to a pH 5.5 stimulus with (red) or without (black) Zn^2+^ preexposure (1 mM, 16 s) recorded from HEK293 cells expressing either wildtype OTOP channels or chimeric channels as indicated. V_m_ was held at −80 mV. The wildtype traces are the same set as shown in Figure 3A. **C.** Average data showing the fold-potentiation after Zn^2+^ pre-exposure (1mM, 16 s) measured from experiments as in (A) and (B). mOTOP2 and its chimeras are shown in blue, mOTOP3 and its chimeras are shown in red. Statistical significance determined using the Krustal-Wallis (non-parametric) test corrected for multiple comparison *, P<0.05. **D, E.** Same data as in (C) plotted to show current magnitudes before and after Zn^2+^ for WT channels and each of the chimeras. **F.** Representative traces of O2/O3(L11-12) currents in response to pH 5.5 after pre-exposure to Zn^2+^ for varying times as indicated. **G.** Average data for experiments as in (F) showing the time-dependence of the potentiation by Zn^2+^ for the O2/O3(L11-12) chimera as compared with WT (n= 4-5). Data of the wildtype OTOP3 are the same set as shown in Figure 2C, D. **H.** Left panel: response of mOTOP2 to alkaline stimulus (pH 9.0) with (red) and without (black) Zn^2+^ pre-exposure (1 mM, 16 s). Right panel: magnitude of currents at pH 9 for WT and mutant channels with and without Zn^2+^-pre-exposure. Currents were smaller after Zn^2+^-pre-exposure for both.

Strikingly, we found that Zn^2+^ potentiation was completely eliminated in a chimera containing the mOTOP3 backbone with the 11-12 linker from mOTOP2 (O3/O2(L11-12); Figure 6 B, C, E). Indeed, even a 64 s exposure to 1mM Zn^2+^ had no effect on the magnitude or activation kinetics of the currents (Supplementary Figure 1). Interestingly, the O3/O2(L11-12) chimera is activated at a higher pH and currents are more rapidly activating than currents carried by mOTOP3 (Figure 6B and (Teng et al., 2022). This suggests that the O3/O2(L11-12) channels may be partly locked in a potentiated state. None of the other chimeras with a mOTOP3 backbone showed a loss of potentiation.

Remarkably, simply transplanting the 11-12 linker from mOTOP3 onto mOTOP2 (O2/O3(L11-12)) conferred sensitivity to potentiation by Zn^2+^. Notably, pre-exposure of O2/O3(L11-12) channels to 1 mM Zn^2+^ for 16s caused a nearly 4-fold potentiation of the subsequent currents evoked in response to a pH 5.5. stimulus (Figure 6A, C, D). The other five chimeras containing an mOTOP2 backbone remained resistant to Zn^2+^ potentiation. Zn^2+^ potentiation of O2/O3(L11-12) showed a time-dependence similar to that of wildtype mOTOP3 channels (Figure 6F, G). mOTOP2 and O2/O3(L11-12), but not mOTOP3, carry outward currents in response to alkaline stimuli (Teng et al., 2022). Thus, we wondered if the sensitivity to potentiation by Zn^2+^ would be observed for outward currents. Following exposure to 1 mM Zn^2+^ for 16s, currents elicited in response to a pH 9.0 stimulus were smaller than in the absence of Zn^2+^ for both WT and O2/O3(L11-12) channels (Figure 6H). We conclude that the 11-12 linker contributes to the potentiation of OTOP channels by Zn^2+^ in response to acidic but not alkaline stimuli.

While none of the other chimeras containing an mOTOP3 backbone showed a reduction in Zn^2+^ potentiation, several showed an increase (Fig 6C, E, and Fig 6). Interestingly, the O3/O2(L1-2) chimera which was previously observed to be non-conductive in response to changes in extracellular pH (Teng et al., 2022), was strongly activated by Zn^2+^. Thus, for this channel, the increase in fold-potentiation mostly reflects the large decrease in the acid-induced currents, rather than an increase in the magnitude of the currents after Zn^2+^ exposure and suggests that the 1-2 linker plays a role in acid-activation of mOTOP3.

To determine if potentiating and inhibiting effects of Zn^2+^ were mediated by the same binding site, we next determined if O3/O2(L11-12) and other mOTOP3 chimeras retained sensitivity to inhibition by Zn^2+^ (Figure 7). All chimeras were inhibited by 1 mM Zn^2+^, including the O3/O2(L11-12) chimera, although the extent of the inhibition, which ranged from ~40% to 80%, varied significantly between some of the chimeras and WT channels (Figure 7). Interestingly, potentiation following Zn^2+^ inhibition was absent not just in the O3/O2(L11-12) chimera, as expected, but also in the O3/O2(L3-4) chimera which showed potentiation with the pre-exposure protocol. This suggests that while differences in L3-4 do not account for differences in Zn^2+^potentiation between mOTOP2 and mOTOP3, residues in L3-4 may nonetheless contribute to potentiation.

**Figure 7.**
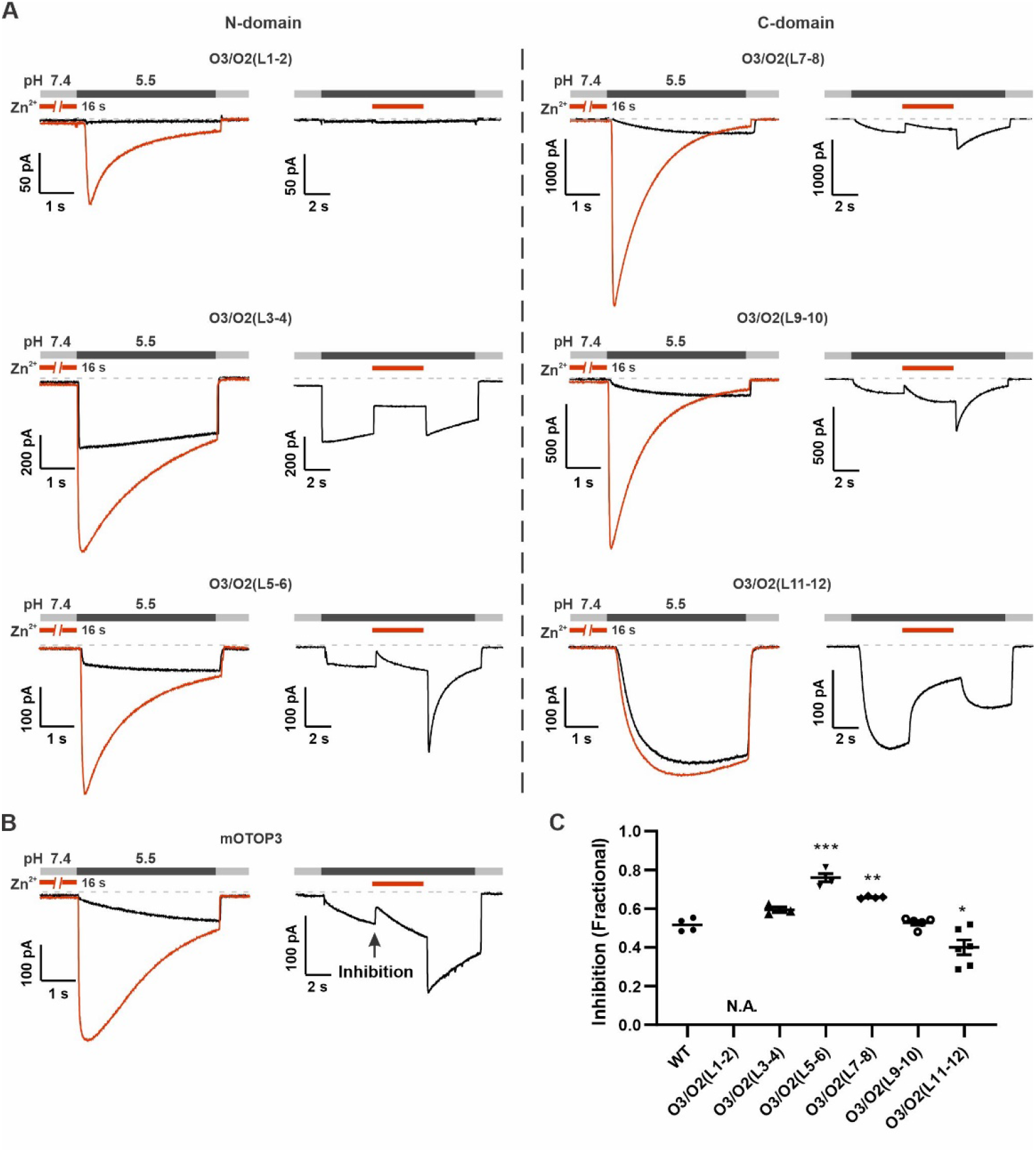
Inhibition of mOTOP3 by Zn^2+^ is retained in chimeric channels. A. Zn^2+^ sensitivity of chimeric mOTOP3-mOTOP2 channels as measured with a pre-exposure protocol (left panel in each) or by adding 1 mM Zn^2+^ to the pH 5.5 stimulus (blocking protocol; right panel in each). Chimeras containing mOTOP2 N-domain and C-domain linkers are shown in the left column, and right columns, respectively. Data from pre-exposure experiments is also presented in Figure 6, and here is shown for comparison to results with the blocking protocol. B. Representative traces of wildtype OTOP3 currents in response to the same protocols as in (A). The arrow indicates the time point in this trace where inhibition by Zn^2+^ was measured, by comparison with the current magnitude before adding Zn^2+^. **C**. Average (mean +/− s.e.m.) and all points data showing fractional inhibition of currents by 1 mM Zn^2+^ measured from wildtype OTOP3 and its chimeras. All channels were similarly inhibited by 1 mM Zn^2+^. Significance determined by ANOVA; *** P<0.0001, ** P=0.005, * P=0.01.

Together we conclude that the mOTOP3 11-12 linker is both necessary and sufficient to confer sensitivity to potentiation by Zn^2+^. Other extracellular linkers may tune the potentiating effects of Zn^2+^, either directly or through allosteric effects on the gating of the channels.

### H531 in the 11-12 linker is critical for Zn^2+^ potentiation

The 11-12 linker is relatively short, consisting of sixteen amino acids, of which five are conserved between the three murine OTOP channels (Figure 8A). In this region, we identified residues that could coordinate Zn^2+^ and that vary between mOTOP3 and mOTOP2 as H531, E533 and E535 (mOTOP3 numbering, Figure 8A). Within mOTOP3 we mutated each residue to that found in mOTOP2 or to alanine. Strikingly, mutation of H531 to either arginine (found in mOTOP2) or alanine eliminated the ability of Zn^2+^ to potentiate mOTOP3 currents, assessed either by measuring the magnitude of the currents or their activation kinetics (Figure 8B – D). In contrast, mutation of the acidic residues (E533 and E535) had more subtle effects: mutation to residues found in mOTOP2 (H and S respectively) had no effect on potentiation while mutation of E535 to alanine significantly reduced, but did not eliminate, potentiation (Figure 8E – H). Interestingly, the corresponding residue in human OTOP1 (D570) was recently reported to regulate proton permeation based on cysteine scanning mutagenesis (Li et al., 2022). Together these results suggest that H531 forms part of the Zn^2+^ binding site.

**Figure 8.**
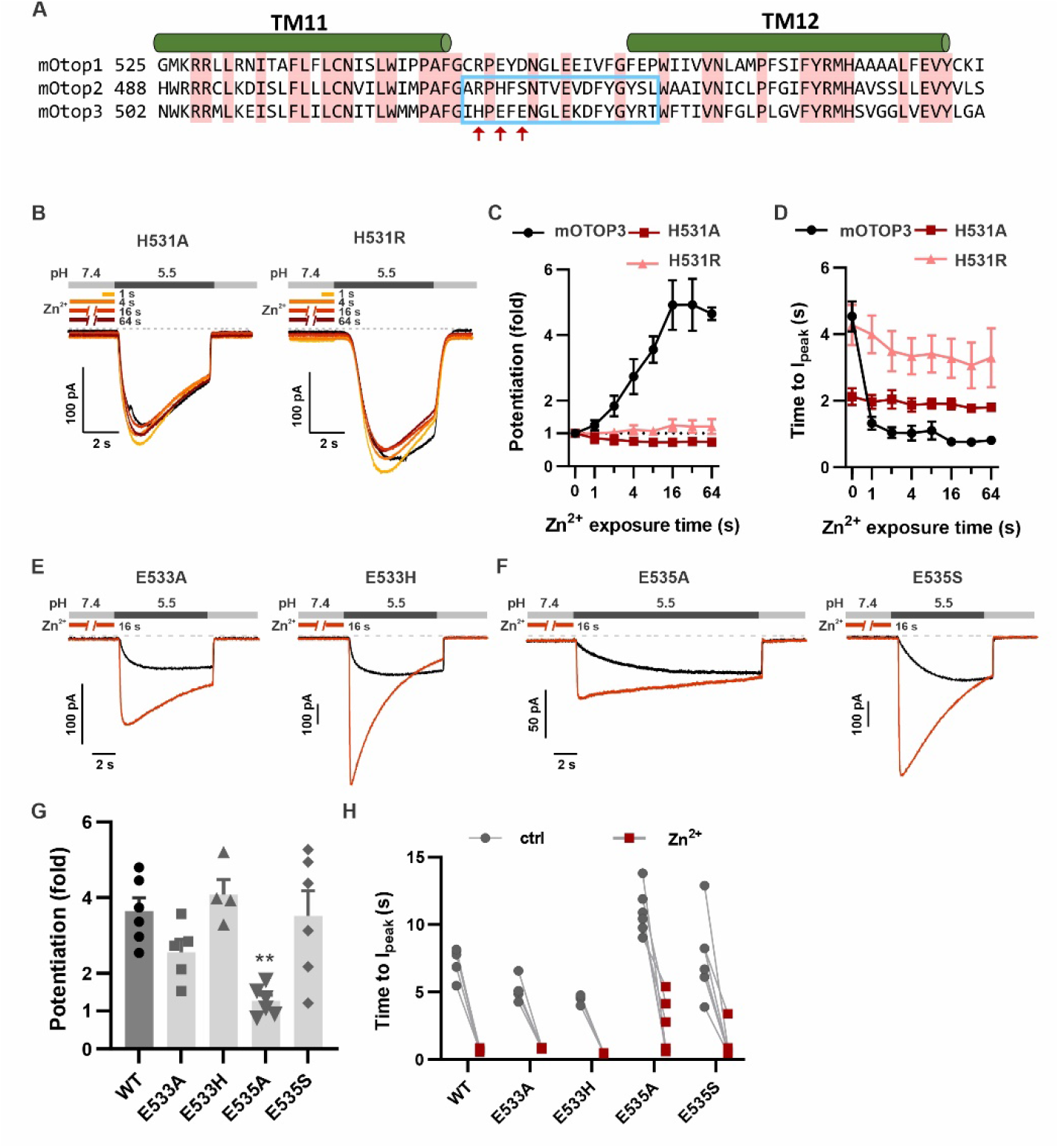
H531 in mOTOP3 L11-12 is essential for Zn^2+^ potentiation. **A.** Sequence alignment of three mOTOP channels. The residues that were exchanged between mOTOP2 and mOTOP3 in the L11-12 chimeras is indicated with a blue box. Residues that differed between the two channels and that were tested are indicated by red arrows. **B**. Representative traces of OTOP3_H531A and H531R currents in response to pH 5.5 after preexposure to Zn^2+^ for varying times as indicated. **C, D**. Average data for fold potentiation (C) and latency to I_peak_ (D), measured from experiments as in B, plotted as a function of pre-exposure time to Zn^2+^ (n= 3 – 7). Data from wildtype OTOP3 are the same set as shown in Figure 2C, D. **E, F.** Responses of OTOP3 mutants as indicated in response to pH 5.5 with (red) or without (black) Zn^2+^ pre-exposure (1mM, 16 s). V_m_ was held at −80 mV. **G.** Average data for fold potentiation measured from experiments as in E and F. Statistical significance determined using the Krustal-Wallis (non-parametric) test corrected for multiple comparison. **, P<0.01. **H.** Latency to peak currents of wildtype OTOP3 or mutant currents, measured from experiments as in E and F. Each set of points represents a separate cell.

### Kinetic model for Zn^2+^ potentiation

These data suggest that Zn^2+^ may act to lock the channel in an open state, possibly by binding more strongly to the same site that is titrated by H^+^ ions to activate the channel (Teng et al., 2022). The data also suggest that a separate site with a faster off-rate mediates inhibition by Zn^2+^. Thus, in the presence of Zn^2+^, the channels may enter a state that is simultaneously activated (open gate) and inhibited. Upon removal of Zn^2+^, the channels may then transit through a fully open state, as the inhibition is relieved faster than the activation. To formally test these predictions, we generated a kinetic model reflecting these properties and asked if it could recapitulate our experimental observations.

We postulated a model comprised of interacting elements that can each transition between two configurations: a pore gate that is either closed or open, a binding site for protons, and two types of binding sites for Zn^2+^ (one activating and one blocking), that are either unoccupied or occupied (Figure 9A). For a detailed description of this type of model representation see Goldschen-Ohm et al., 2014. The proton and activating Zn^2+^ sites are energetically coupled to the pore such that binding speeds pore opening and/or slows pore closing. To model the competition of protons and Zn^2+^ for the same activating site(s), we destabilized all states where both the proton and activating Zn^2+^ sites are simultaneously occupied. We modeled the blocking Zn^2+^ site as independent of all other model elements, except that when occupied all channel current is blocked. Simulated currents in response to the same pH and Zn^2+^ protocols used in our experiments qualitatively describe our observed mOTOP3 current responses (Fig. 9B, C), supporting the plausibility of our proposed mechanism. For example, the model recapitulates the rebound following addition and removal of Zn^2+^ during an acid stimulus. It also recapitulates the speeding up of current activation for a pre-exposure to Zn^2+^ at 0.3 mM, and the increase in current magnitudes with pre-exposure to Zn^2+^ at 1 and 3 mM.

**Figure 9.**
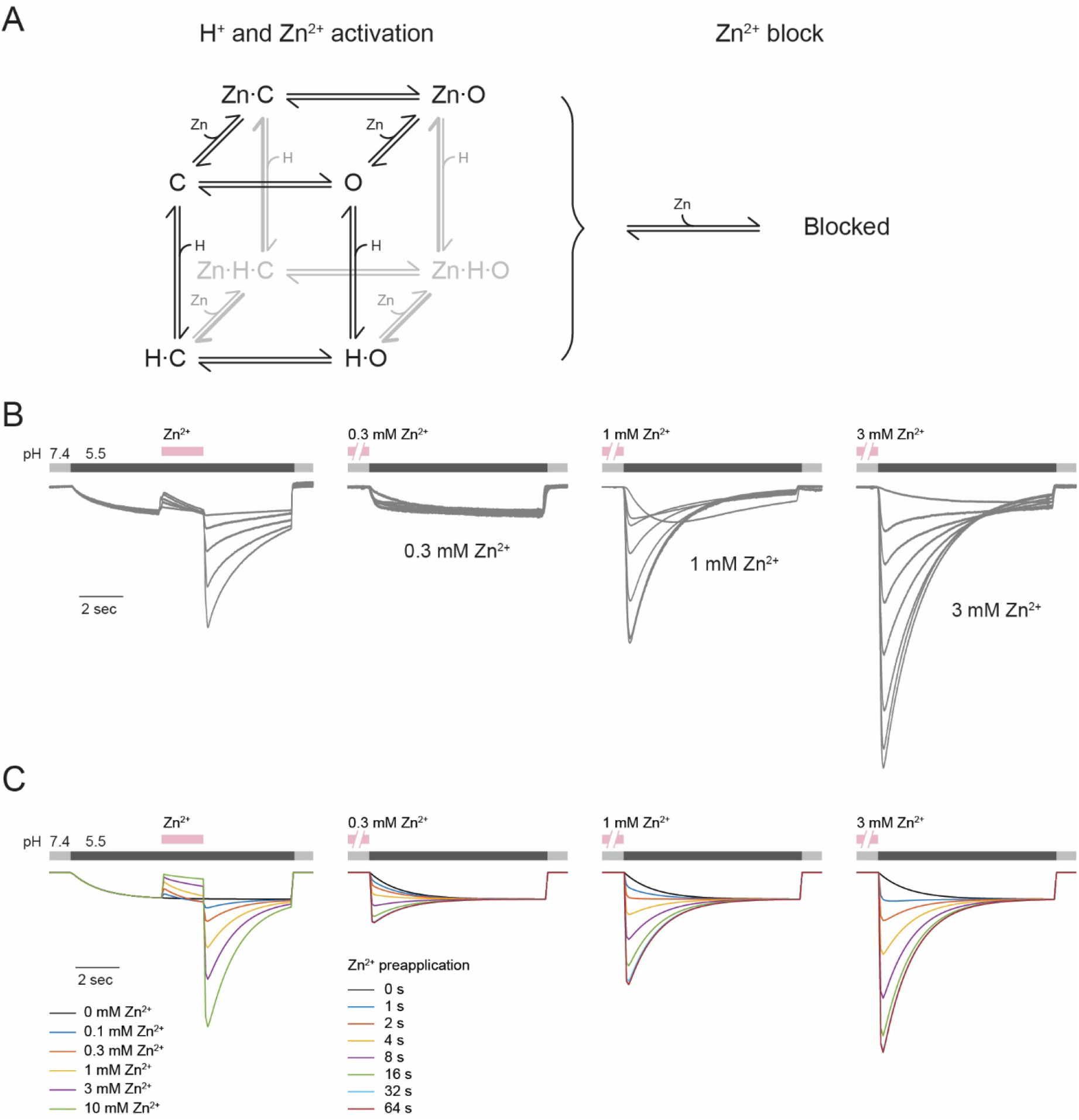
OTOP3 kinetic model for Zn^2+^ potentiation and block. **A.** Kinetic model for activation of OTOP3 by H+ and Zn2+. The channel moves from a closed state (C) upon binding H+ or Zn2+ to an open state (Zn-O, H-O) in which it permeates protons. The doubly bound Zn-H-O and Zn-H-C states are disfavored energetically. A separate site binds Zn2+ and inhibits channel permeation, independently of the gating state. **B**. Example OTOP3 current recordings from Figs. 1C and Fig. 2B. Each set of recordings was from a different cell. **C.** Simulated currents for the model under the same protocols as in B. Model parameter were adjusted manually and were the same for all traces. The model replicates the rebound seen upon addition of Zn^2+^ at pH 5.5, and the potentiation seen with pre-exposure to Zn^2+^ at pH 7.4

However, this model also undoubtedly represents an oversimplification of the true number and properties of the states the channel adopts. For example, the model does not recapitulate the observed decay of the currents below baseline when strongly potentiated, which may reflect an inactivation process not modeled. Making things more complicated, and interesting, it is also entirely possible that there is more than one permeation pathway and gate, and that Zn^2+^ opens one of the permeation pathways and not the other. The 11-12 loop sits close to the intrasubunit interface, so that at present it is not possible to predict if it would open a permeation pathway in the N domain, the C domain, or the intrasubunit interface.

## Discussion

The OTOP proton channels were discovered in 2018 when OTOP1 was cloned from taste tissue as a putative sour receptor (Tu et al., 2018). At that time, OTOP1 was shown to have near-perfect selectivity for protons over other ions, and to generate inward currents as the pH was lowered. Vertebrate OTOP2 and OTOP3, as well as invertebrate channels also carry inward proton currents in response to extracellular acidification, and where tested have been shown to function as proton-selective ion channels (Tu et al., 2018; Chen et al., 2019) (see also (Teng et al., 2022)). Nearly all descriptions regarding the functional properties of OTOP channels come from the heterologous expression of OTOP channels and save for the description of proton currents now attributed to OTOP1 channels in taste cells (Chang et al., 2010; Bushman et al., 2015), all descriptions of OTOP channels postdate the discovery of the genes encoding the proton channels. That is, even by scouring the literature, it is difficult to find a description of a proton current that could be attributed to an OTOP channel in a native cell type. Thus, in contrast to K^+^channels where there was a vast literature regarding their functional properties before their cloning, for OTOP channels, this kind of information was not available.

Critically, before this work and that described in (Teng et al., 2022), it was not known if OTOP channels were gated. Here we provide evidence that indeed they are. Importantly, we can see a change in the current magnitude under conditions where there is no change in the concentration of the permeant ion (H^+^) or driving force for its entry. In the apo, Zn^2+^-free condition, mOTOP3 currents are small and slowly activating. With pre-exposure to 1-3 mM Zn^2+^ for several seconds, the currents increase in magnitude by up to 10-fold. This can only be explained if Zn^2+^ is either increasing the probability that the channels open or increasing their conductance – gating them. We also find that pre-exposure to copper (Cu^2+^) strongly potentiates OTOP3 currents. Thus, we can say now with certainty that some OTOP channels are gated, and that Zn^2+^ and other transition metals act as positive allosteric modulators of gating.

### Structural considerations

The structures of two of the three OTOP channels have been reported at near-atomic resolution (zebrafish OTOP1, chicken OTOP3, and Xenopus OTOP3; (Chen et al., 2019; Saotome et al., 2019)), and more recently the structures predicted for mouse and human OTOP channels that appear to be reliable (Jumper et al., 2021; Teng et al., 2022; Varadi et al., 2022). All the structures to date share common features: The protomers assemble as dimers, and each protomer shows a two-fold symmetry, leading to an overall pseudo-tetrameric stoichiometry. The four pseudo-subunits come together to form a central cavity that is filled with lipids and cannot support ion permeation. Instead, ions may permeate through the barrel-like structures formed from transmembrane domains 2-6 (N domain) and 8-12 (C domain) or at the intrasubunit interface (formed mostly by tm 6 and 12). From these static structures, it is not possible to tell if the channels are gated, and if so, what state they are in (open or closed). Based on the pH sensitivity of the channels (Teng et al., 2022), we presume they are closed.

We have focused on mOTOP2 and mOTOP3, which show the most divergent functional properties. In addition to differences in the effects of Zn^2+^ described here, we have also described differences in pH sensitivity of the two channels (Teng et al., 2022). mOTOP3 is gated by protons, and thus conducts currents only in response to extracellular acidification (<pH 5.5), while OTOP2 is constitutively open and conducts currents over a large pH range, including outward currents when the extracellular solution is alkalinized. The strikingly different functional properties of the two channels, but otherwise overall similar architecture, provided us the opportunity to identify motifs involved in gating using a chimeric approach.

Remarkably, we found that a short stretch of amino acids linking transmembrane domains 11-12 was necessary for Zn^2+^ potentiation of mOTOP3 and sufficient to confer Zn^2+^-sensitive gating on mOTOP2. Within that stretch, we identified one amino acid, H531, mutations of which (to R or A) completely abolished Zn^2+^-sensitive gating in mOTOP3. As histidine is well documented to form part of the Zn^2+^ binding site in other Zn^2+^-sensitive proteins (Vallee and Auld, 1990), including the voltage-gated proton channel HV1 (Ramsey et al., 2006), H531 likely contributes to coordinating Zn^2+^ in mOTOP3. However, Zn^2+^ is typically coordinated by sidechains of four or more residues, which in addition to the imidazole rings of histidine, includes sulfhydryl groups of cysteine, carboxyl group of acidic residues (aspartic and glutamic acid) in enzymatically active Zn^2+^ binding sites. Moreover, water can also help to coordinate Zn^2+^ (Vallee and Auld, 1990). These other residues may include ones conserved between mOTOP3 and mOTOP2 and need not be on the 11-12 linker. It is tempting to speculate that the binding site spans parts of the channel that are at a distance in the closed state and that come together in the open state as shown for other ion channels (Gordon and Zagotta, 1995b; Zhang et al., 2009). This mechanism would in essence stabilize the open state.

### Role of Zinc in health and disease

Zinc is required for the functioning of a wide range of enzymes and proteins in the body and supplementary zinc has both beneficial as well as detrimental health effects. Given that it is mostly not known which cells express OTOP channels, or what functions they play in those cells, it is hard to know what functional role Zn^2+^-sensitive gating might play in physiology. Meanwhile, the dual blocking and potentiating effect of zinc make it likely that the enhanced activity would only be evident under conditions where zinc concentrations dropped rapidly or that favored binding of Zn^2+^ to its activating site over its blocking site. These studies also raise the interesting prospect that other molecules could be found that function as positive allosteric modulators of OTOP channels, with selectivity for the activating site over the blocking site, and that could ameliorate effects of reductions in OTOP channel function that lead to vestibular or other disorders.

## Materials and Methods

### Clones, cell lines, and transfection

Mouse OTOP1, OTOP2 and OTOP3 cDNAs were in a pcDNA3.1 vector with an N-terminal fusion tag consisting of an octahistidine tag followed by eGFP, a Gly-Thr-Gly-Thr linker and 3C protease cleavage site (LEVLFQGP) were as previously described (Saotome, 2019 #186). All chimeras and mutations were generated using In-Fusion Cloning (Takara Bio) and were confirmed by Sanger sequencing (Genewiz) (see (Teng et al., 2022)).

HEK293 cells were purchased from ATCC (CRL-1573). The cells were cultured in a humidified incubator at 37 °C in 5% CO_2_ and 95% O_2_. The high glucose DMEM media (ThermoFisher) is implemented with 10% fetal bovine serum (Life Technology) and 1% Penicillin-streptomycin antibiotics. Cells were passaged every 3 – 4 days.

Cells used for patch-clamp recordings were transfected in 35mm Petri dishes, with 600 – 1000 ng DNA and 2 μL TransIT-LT1 transfection reagents (Mirus Bio Corporation) following the manufacturer’s protocol. After 24 hours, the cells were lifted using Trypsin-EDTA and plated into a recording chamber.

### Patch-clamp electrophysiology

Whole-cell patch-clamp recording was performed as previously described (Teng 2019). Briefly, recordings were obtained with an Axonpatch 200B amplifier, digitized with a Digidata 1322a 16-bit data acquisition system, acquired with pClamp 8.2, and analyzed with Clampfit 9 or 10 (Molecular Devices). Records were sampled at 5 kHz and filtered at 1 kHz. Patch pipettes with a resistance of 2 – 4 MΩ were fabricated from borosilicate glass (Sutter instrument). Solution exchange was achieved with a fast-step perfusion system (Warner instrument, SF-77B) custom modified to hold seven microcapillary tubes in a linear array. The membrane potentials were held at −80 mV unless otherwise indicated.

### Patch-clamp electrophysiology solutions

Tyrode’s solution contained 145 mM NaCl, 5 mM KCl, 1 mM MgCl2, 2 mM CaCl2, 20mM dextrose, 10mM HEPES (pH adjusted to 7.4 with NaOH). Standard pipette solution contained 120 mM Cs-aspartate, 15 mM CsCl, 2 mM Mg-ATP, 5 mM EGTA, 2.4 mM CaCl2 (100nM free Ca2+), and 10 mM HEPES (pH adjusted to 7.3 with CsOH; 290 mOsm). Standard Na+ free extracellular solutions contained 160 mM NMDG, 2 mM CaCl2, and 10 mM buffer (HEPES for pH 7.4, MES for pH 6.0 and 5.5), and pH was adjusted with HCl to 7.4.

ZnCl_2_ was directly introduced into the Na^+^-free solutions where the final concentration was less than 3 mM, which caused a change in osmolarity of <10 mOsm. The pH was re-adjusted with NMDG-OH or HCl as needed. For the experiment in Figure 1C, the concentration of NMDG in the 10 mM Zn^2+^ solution was reduced to maintain a ~300 mOSM osmolarity.

For experiments in figure 5, FeCl_2_, CoCl_2_, NiCl_2_, CuCl_2_, and CdCl_2_ were directly introduced into the Na^+^-free solutions at a concentration of 1 mM immediately before the experiment was performed. The pH was re-adjusted with NMDG-OH or HCl as needed.

### Quantification and statistical analysis

All data are presented as mean ± SEM if not otherwise noted. Statistical analysis was performed using Graphpad Prism 9 (Graphpad Software Inc). The sample sizes in this study are similar to those in the literature for similar studies. For comparison of fold-potentiation, the Krustal-Wallis (non-parametric) test was used, with correction where appropriate for multiple comparison using the two-stage linear step-up procedure of Benjamini, Krieger and Yekutieli (Graphpad Prism 9). All replicates are biological replicates. Number of replicates are indicated in the figure or figure legend. They represent recordings from different cells transfected with the same plasmid DNA. Data were excluded if a gigaohm seal was not established or maintained, as indicated by an inward current of >80 pA (−80 mV) in the presence of the OTOP channel blocker Zn2+ applied at pH 7.4. Data sets that represent time series were excluded if four or fewer time points were collected out of a possible eight time points, due to seal instability.

Representative electrophysiology traces shown in the figures were acquired with pClamp, and in some cases, the data was decimated by 10-fold before exporting into graphic programs, Origin (Microcal) and Coreldraw (Corel).

### Kinetic Modeling

A kinetic model was generated based on interacting binary elements as described in (Goldschen-Ohm et al., 2014). Briefly, each element can adopt one of two possible configurations (closed or open for the pore, unbound or bound for each binding site) with intrinsic rate constants for transitioning between them. The elements are energetically coupled such that binding at a particular site either promotes or inhibits pore opening. To simulate competition between H^+^ and Zn^2+^ for the activating site(s), occupation of one site energetically destabilizes the other.

Model parameters were manually adjusted to qualitatively recapitulate the experimental observations. Rate constants for pore opening/closing (*k_o_/k_c_*), proton binding/unbinding (*k_onH_/k_offH_*), Zn^2+^ binding/unbinding to activating sites (*k_onP_/k_offP_*) and blocking site (*k_onB_/k_offB_*) are (s^-1^ or M^-1^s^-1^ for binding rates): *k_o_* = 0.02, *k_c_* = 25, *k_onH_* = 5×10^4^, *k_offH_* = 1, *k_onP_* = 35, *k_offP_* = 1, *k_onB_* = 5×10^4^, and *k_offB_* = 20. The interaction between the proton site and pore gate is such that proton binding reduces the energy barrier for pore opening by −4 kcal/mol. Reciprocally, pore opening reduces the energy barrier for proton binding by −1 kcal/mol, and increases the barrier for proton unbinding by 3 kcal/mol. To simulate competition between protons and Zn^2+^ for the activating site, states with both sites occupied were destabilized by 6 kcal/mol. Zn^2+^ binding to each activating site reduces the energy barrier for pore opening by −3 kcal/mol for both Models 1 and increases the barrier for pore closure by 3 kcal/mol for Models 1. Reciprocally, pore opening reduces the barrier for Zn^2+^ binding to each activating site by −5 kcal/mol and increases the barrier for Zn^2+^ unbinding from each activating site by 1 kcal. The Zn^2+^ blocking site was assumed to be independent of all other elements, but when occupied blocks all channel current.

For comparison with model simulations, the entire set of currents across all cells was uniformly scaled under the assumption that the maximal response following preapplication of 3 mM Zn^2+^ is reflective of channels with an open probability of ~0.9, a measure that has not been experimentally verified. It is likely that the same model structure will similarly describe responses with lower maximal open probabilities, albeit with slightly different parameter values. Given these uncertainties, we did not attempt to optimize the model parameters, but instead manually identified parameters that qualitatively recapitulate the data. The time-dependent probability in each state was simulated as described by (Colquhoun and Hawkes, 1995) after first generating the matrix of transition rates between states from the model’s binary elements and interactions(Goldschen-Ohm et al., 2014). Currents were computed from the simulated open probability assuming a conductance of 1 pS and a driving force of −80 mV in pH 5.5 and zero in pH 7.4 as estimated from ramp experiments (Teng et al, 2022a). The choice of conductance is arbitrary given that we do not know how many channels are in each recording, and thus only contributes to the overall scaling of the current.

## Acknowledgments

We thank Jackson Walker, Kevin Chyung and Anne Tran for expert technical support and all members of the Liman and Ward labs for helpful discussions. We also thank Z. Qui for generously sharing cell lines. This research was supported by NIH grants R01DC013741 and R01GM131234 to E.R.L.

## Material availability

All materials generated during this study including mutant channels are available upon request.

## Data availability

All data generated or analyzed during this study are included in the manuscript and supporting files. The source code for simulations is provided in supplementary information.

## Competing interests

The authors declare no competing interests

---

**Supplementary Figure 1.**
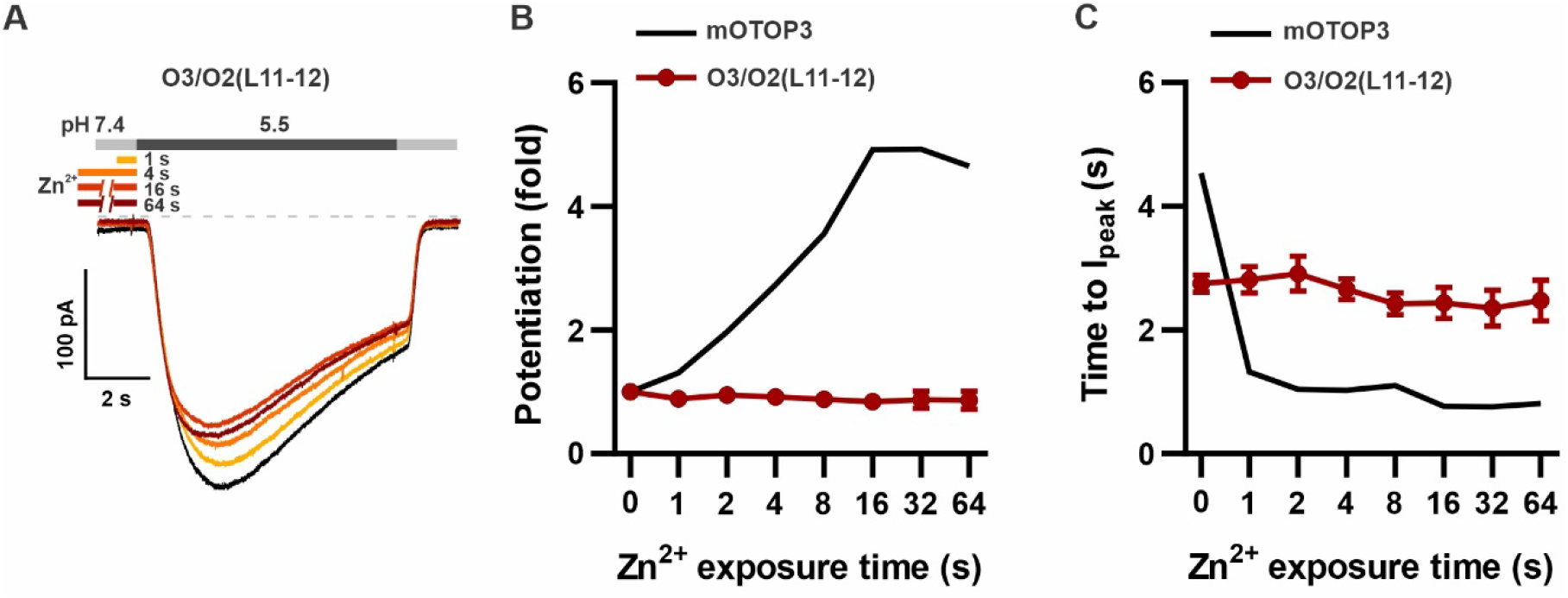
O3/O2(L11-12) chimera is completely insensitive to potentiation by Zn^2+^. **A.** Representative traces show O3/O2(L11-12) currents in the absence of Zn^2+^ (black) and with pre-exposure times as indicated. **B,C**, Average data for peak (B) currents and time to peak (C) O3/O2L(11-12) currents as in (A), or mOTOP3 channels (data for the wildtype OTOP3 are the same as shown in Figure 2C, D) (n=4-5).

## Notes

### Competing Interest Statement

The authors have declared no competing interest.

